# Inherent Metabolic Adaptations in Adult Spiny Mouse (*Acomys*) Cardiomyocytes Facilitate Enhanced Cardiac Recovery Following Myocardial Infarction

**DOI:** 10.1101/2024.05.22.595229

**Authors:** Annapurna Kuppa, Afnan Alzamrooni, Rachel Lopez, Tahra Suhan, Rajesh Chaudhary, Nicole Collins, Fran Van den Bergh, Riham Abouleisa, Harrison Wang, Tamer Mohamed, Jonathan Satin, Costas Lyssiotis, Daniel A. Beard, Ahmed Abdel-Latif

## Abstract

The adult mammalian heart has limited regenerative capacity following injury, leading to progressive heart failure and mortality. Recent studies have identified the spiny mouse (*Acomys*) as a unique model for mammalian cardiac isch3emic resilience, exhibiting enhanced recovery after myocardial infarction (MI) compared to commonly used laboratory mouse strains. However, the underlying cellular and molecular mechanisms behind this unique response remain poorly understood. In this study, we comprehensively characterized the metabolic characteristics of cardiomyocytes in *Acomys* compared to the non-regenerative *Mus musculus*.

We utilized single-nucleus RNA sequencing (snRNA-seq) in sham-operated animals and 1, 3, and 7 days post-myocardial infarction to investigate cardiomyocytes’ transcriptomic and metabolomic profiles in response to myocardial infarction. Complementary targeted metabolomics, stable isotope-resolved metabolomics, and functional mitochondrial assays were performed on heart tissues from both species to validate the transcriptomic findings and elucidate the metabolic adaptations in cardiomyocytes following ischemic injury.

Transcriptomic analysis revealed that *Acomys* cardiomyocytes inherently upregulate genes associated with glycolysis, the pentose phosphate pathway, and glutathione metabolism while downregulating genes involved in oxidative phosphorylation (OXPHOS). These metabolic characteristics are linked to decreased reactive oxygen species (ROS) production and increased antioxidant capacity. Our targeted metabolomic studies in heart tissue corroborated these findings, showing a shift from fatty acid oxidation to glycolysis and ancillary biosynthetic pathways in *Acomys* at baseline with adaptive changes post-MI. Functional mitochondrial studies indicated a higher reliance on glycolysis in *Acomys* compared to *Mus*, underscoring the unique metabolic phenotype of *Acomys* hearts. Stable isotope tracing experiments confirmed a shift in glucose utilization from oxidative phosphorylation in *Acomys*.

In conclusion, our study identifies unique metabolic characteristics of *Acomys* cardiomyocytes that contribute to their enhanced ischemic resilience following myocardial infarction. These findings provide novel insights into the role of metabolism in regulating cardiac repair in adult mammals. Our work highlights the importance of inherent and adaptive metabolic flexibility in determining cardiomyocyte ischemic responses and establishes *Acomys* as a valuable model for studying cardiac ischemic resilience in adult mammals.

**Graphical abstract:** 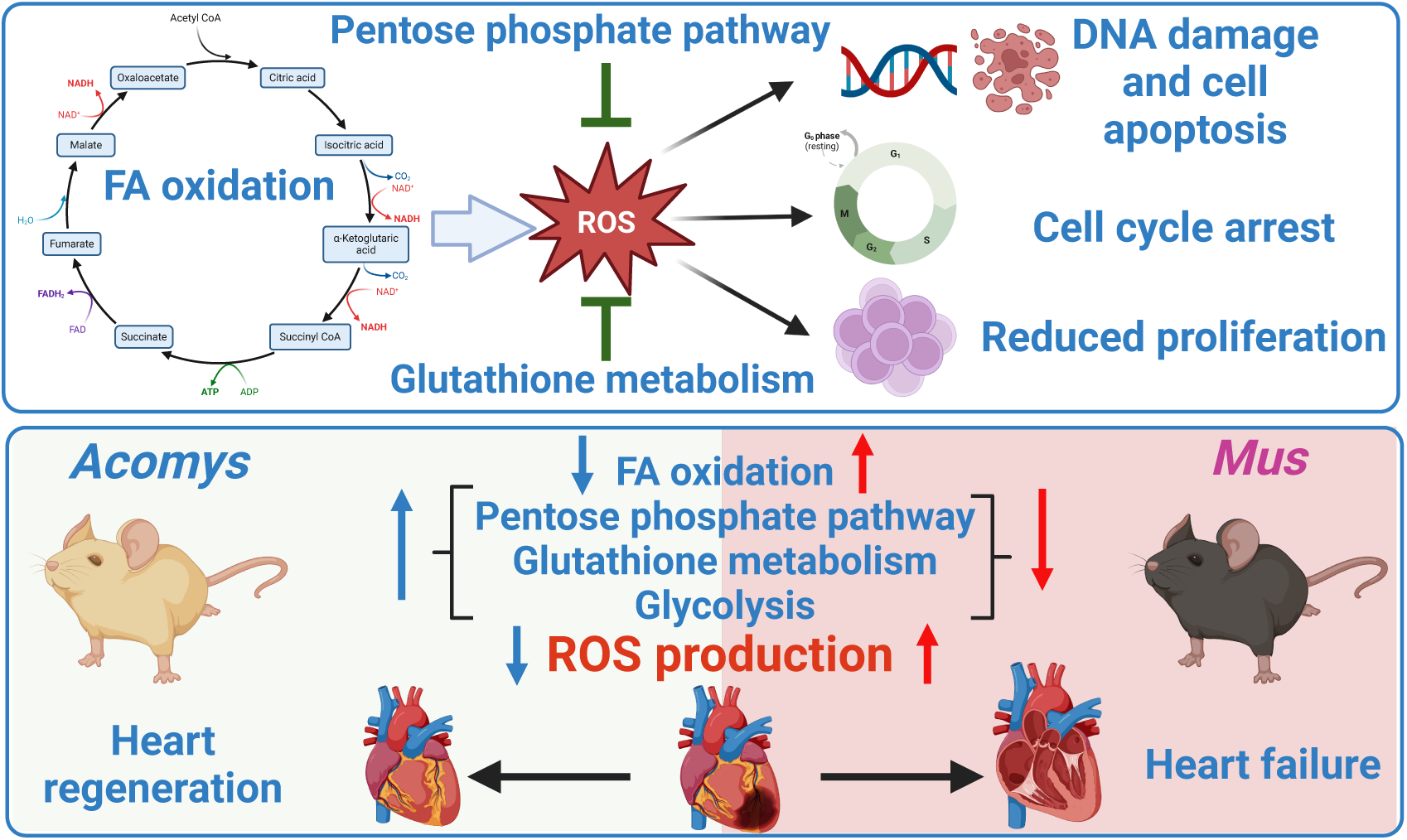

## Introduction

Myocardial infarction (MI) is a leading cause of morbidity and mortality worldwide, resulting in the loss of functional cardiomyocytes, cardiac remodeling, and ultimately heart failure [1]. In adult mammals, the heart has limited regenerative capacity, and the primary response to injury is fibrotic repair [2]. In contrast, the hearts of neonatal mammals and adult zebrafish can efficiently regenerate after injury through the proliferation of pre-existing cardiomyocytes [3–6]. However, these models do not accurately represent the complex pathophysiology in adult mammals. As a result, new strategies to preserve cardiac tissue and improve heart function in treating ischemic heart disease remain an unmet clinical need. The emergence of spiny mice (*Acomys*) as an adult mammalian model with enhanced cardiac repair capabilities [7] may help bridge the gap between regenerative biology and clinical medicine, offering innovative solutions to protect the heart after myocardial infarction (MI). Elucidating the mechanisms underlying successful cardiac recovery in *Acomys* can provide valuable insights for developing novel therapies to promote cardiac repair in adult mammals.

Recent studies have highlighted the critical role of metabolism in regulating cardiomyocyte proliferation and heart regeneration. In zebrafish, cardiomyocytes in the regenerating area undergo a metabolic shift from oxidative phosphorylation to glycolysis, which is essential for their proliferation [8]. Similarly, in neonatal mammals, glycolysis is the predominant form of metabolism and supersedes fatty acid oxidation. However, a metabolic switch from glycolysis to fatty acid oxidation occurs in the first week after birth, linked to loss of regenerative ability in mammals. This metabolic switch has been linked to increased levels of intracellular reactive oxygen species (ROS) in CMs, resulting in DNA damage and cessation of proliferation [9, 10]. As such, strategies targeting metabolic pathways to reduce ROS production have been shown to reactivate the ability of CMs to proliferate and enhance cardiac regeneration in adult mammals [11, 12]. Animal studies showed that inhibiting glycolysis impairs neonatal heart regeneration while activating glycolysis enhances cardiomyocyte proliferation and promotes regeneration in adult mice [13]. Furthermore, the Hippo/YAP signaling pathway has also been implicated in regulating cardiomyocyte proliferation and heart regeneration. Activating YAP in adult mouse hearts promotes cardiomyocyte dedifferentiation and proliferation, which involves a partial reprogramming towards a neonatal-like state [14]. This reprogramming is associated with metabolic changes, including increased glycolysis and decreased oxidative phosphorylation [14], further supporting the link between metabolism and cardiomyocyte proliferation.

Further evidence for the importance of metabolism in cardiomyocyte proliferation comes from studies using cell cycle regulators to induce cell division. In adult mouse cardiomyocytes and human induced pluripotent stem cell-derived cardiomyocytes (hiPS-CMs), overexpression of cell cycle regulators (4F: *Cdk1*, *Cdk4*, *Ccnb*, *Ccnd*) downregulates oxidative phosphorylation genes and upregulates genes involved in glycolysis and ancillary biosynthetic pathways [15]. Metabolic analyses revealed that 4F expression decreases glucose oxidation and augments glucose-derived carbon allocation into the NAD+, glycogen, hexosamine, phospholipid, and serine biosynthetic pathways [15]. Inhibiting these biosynthetic pathways decreased cell cycle entry while overexpressing *Pck2*, which drives carbon from the Krebs cycle to the glycolytic pool, enhanced cell cycle progression [15].

Unlike most adult mammals, the hearts of *Acomys* (*Acomys dimidiatus*) demonstrate enhanced recovery in response to MI [7, 16]. While *Acomys* have emerged as a novel adult mammalian model to study cardiac injury, little is known about the molecular mechanisms underlying their ischemic resilience response. Therefore, in this study, we investigated the transcriptomic and metabolomic profiles of CMs on sham-operated, 1-, 3-, or 7-days post-MI in adult *Acomys* and *Mus*. We found a distinct CM population that exhibits a shift in metabolism toward glycolysis in *Acomys*, while *Mus* continues to rely on fatty acid oxidation post-injury. Furthermore, our functional analyses corroborated our RNAseq findings. Together, these findings highlight the critical role of *Acomys* CM metabolic phenotypes in Ischemic recovery. *Acomys* hearts exhibit a shift from oxidative metabolism to glycolysis and activation of biosynthetic pathways, which could be essential for their survival. Understanding these metabolic adaptations’ complex and synergistic interplay may provide new targets for enhancing cardiac ischemic recovery in adult mammals.

## Materials and methods

### Animals

All animal work described in the manuscript was carried out in accordance with the University of Michigan Institutional Animal Care and Use Committees (IACUC) under the protocol PRO00010623, and the University of Kentucky IACUC protocol 2019-3254, 2013-1155, 20213739, and 2011-0889. Male *Mus musculus* (sexually mature animals, 8–12 weeks) C57BL6/J (strain #000664) were obtained from the Jackson Laboratory, Bar Harbor, ME. Male *Acomys dimidiatus* (sexually mature animals, 6–8 months old) were obtained from our in-house breeding colony. *Mus* were maintained on a 18% mouse chow (Tekland Global 2918, Harlan Laboratories, Indianapolis, IN) diet and housed in a standard 12h light/dark cycle in a temperature and humidity-controlled room. *Acomys* were maintained on a 14% mouse chow (Teklad Global 2014, Harlan Laboratories) diet along with black sunflower seeds and housed separately in a standard 12h light/dark cycle with exposure to natural light.

### Myocardial Infarction (MI) Surgery

MI surgery was performed as previously described [7]. In brief, we anesthetized mice with 1–3% isoflurane using a small animal vaporizer system. The pain reflex was examined to make sure that the mice were adequately anaesthetized before surgery. A thoracotomy was performed between the 4th and 5th ribs and the pericardial sac was removed. The heart was exposed and pushed out of the chest. The left anterior descending coronary artery (LAD) was identified under direct vision and was permanently ligated 3 mm below its origin using 6–0 silk suture. After LAD ligation, the heart was returned to the intrathoracic space, the muscles were closed, and the skin was sutured using 4–0 proline running sutures.

### EdU Administration

To assess cardiomyocyte proliferation, EdU-positive cardiomyocytes were quantified in both the peri-infarct and remote regions in all strains 17 days post-MI. Mice received a single intraperitoneal injection of 5-ethynyl-2’-deoxyuridine (EdU, 10mg/kg, Cayman Chemical, Ann Arbor, MI) immediately after MI surgery and continued daily through post-op Day 14.

### Tissue collection and Histology

Tissue (heart and lung) from all species were collected and weighed immediately at sacrifice. The dry lung weight was collected after 3 days of 65° C incubation. Hearts were perfused with PBS (VWR International) followed by 4% PFA (VWR International) fixation via cannulation of the ascending aorta. Hearts were post-fixed overnight at 4°C. Hearts were then sectioned in half long axis at the level of the ligation and transferred to 70% ethanol until sectioning. Tissues were then paraffin-embedded and sectioned. DAPI counterstain on ventricular heart sections was performed to analyze the cellular density of scar tissue. Hearts were stained for cardiac troponin T (cTNNT) and EdU with a DAPI counterstain. Images were taken with a 40x oil immersion objective on a Nikon A1 Confocal Microscope in the University of Kentucky Confocal Microscopy facility. 8-15 images per section were taken, and the data reported as a percentage of cTNNT+ cells that also expressed EdU.

### Cardiomyocyte cell fate tracking

To quantify cytokinesis events, we recently developed a cytokinesis/M-phase reporter using an Aurora kinase B (*AurKb*) promoter-based GFP reporter and cloned it into a non-integrating lentivirus to drive the permanent expression of GFP protein specifically in cardiomyocytes (CMs) using the TNNT promoter [17]. *AurKb* is one of the major protein kinases that ensures the proper execution and fidelity of mitosis and is only expressed for a short time during the M-phase/cytokinesis process, localizing to the central spindle during anaphase and in the midbody during cytokinesis [18].

### In vitro studies

*Acomys* neonatal CMs were isolated as previously described [19] and cultured on laminin (Gibco) coated plates. Cells were then transfected with IL DR and IL Aurora cre constructs (2×10^5 TU/ml), and imaged using fluorescent microscopy.

### In vivo studies

To assess CM proliferation in vivo, we delivered the AurKb-GFP-IL lentiviral reporter system to the peri-infarct region at 4.5×10^7 transducing units (TU) intramyocardially immediately following left anterior descending (LAD) coronary artery ligation. The rate of GFP+ cells was assessed at 14 days post-injury (dpi) in frozen sections co-stained with *cTnnt* to provide definitive evidence regarding CM proliferation across species. All measurements were analyzed by blinded observers.

Images of DAPI stained tissue were taken at 4x and 60x magnification using Olympus BX53 microscope in the University of Kentucky Light Microscopy Core. The number of nuclei in each imaging field was counted using ImageJ.

### Mass spectrometry

Sample preparation: samples were extracted using a solvent mixture of methanol, acetonitrile, and acetone (1:1:1) with internal standards. For 200 samples, 100 mL of extraction solvent and 4 mL of internal standard mixture were used. Samples (100 µL) were pipetted into 1.5 mL micro-centrifuge tubes, followed by 400 µL of extraction solvent. The mixture was vortexed for 5 minutes, left to sit at 4°C for 30 minutes, vortexed again, and then left at -20°C for 1 hour. After centrifuging at 15,000 rpm for 10 minutes, 250 µL of the supernatant was transferred to new tubes, dried under a nitrogen stream, and reconstituted in 100 µL of methanol:water (2:98). The samples were vortexed and centrifuged again before transferring the supernatant to autosampler vials.

Agilent Technologies 6530 Accurate-Mass Q-TOF with a dual ASJ ESI ion source was used as the mass detector. Mass spectrometer settings were as follows: Ion source: gas temperature - 325 ℃, drying gas flow - 10 l/min, nebulizer pressure - 45 psig, sheath gas temperature - 400 ℃, sheath gas flow - 12 l/ml, capillary voltage - 4000 V. fragmentor voltage - 140 V, skimmer voltage - 65 V, mass range 50-1000 m/z, acquisition rate 2 spectra/s. Inline mass calibration was performed using debrisoquine sulfate (m/z 176.1182) and HP-0921 from Agilent (m/z 922.0098) in positive mode and 4-NBA (m/z 166.0146) and HP-0921 from Agilent (m/z 966.0007, formate adduct) in negative mode.

### Nuclei Isolation

Cardiomyocyte nuclei isolation was performed as previously described [20] with modifications. All chemicals used were purchased from Sigma Aldrich, unless otherwise indicated. For each nuclear sample, 4-5 tissue samples were collected, pooled together, and minced with a razor blade. Minced samples were resuspended in 5 mL Lysis Buffer (320 mM sucrose, 10 mM Tris-HCl, 5 mM CaCl2, 5 mM Magnesium Acetate, 2 mM EDTA, 0.5 mM EGTA, 1 mM DTT, 1x complete protease inhibitor (Roche), 200U/mL RNase OUT Recombinant Ribonuclease Inhibitor (Invitrogen, 10777–019), pH 8) and homogenized with a Dounce grinder for 15 strokes. The lysate was sequentially filtered through 70 μm and 40 μm cell strainers (Falcon, 352350 and 352340) and centrifuged at 1,000 × g for 5 min at 4 °C to pellet nuclei. The nuclear pellet was subsequently resuspended in 2 mL Sucrose Buffer (1 M sucrose, 10 mM Tris-HCl, 5 mM magnesium acetate, 1mM DTT, 1x complete protease inhibitor, 200 U/mL RNase OUT Recombinant Ribonuclease Inhibitor, pH=8) and the suspension was cushioned on top of 4 mL Sucrose Buffer, followed by centrifugation at 1,000 × g for 5 min at 4 °C to pellet nuclei. The nuclear pellet was subsequently washed once in 1 mL of Nuclei Storage Buffer (NSB) (440 mM sucrose, 10 mM Tris-HCl, 70 mM KCl, 10 mM MgCl2, 1.5 mM spermine, 1x complete protease inhibitor, 200 U/mL RNase OUT Recombinant Ribonuclease Inhibitor, pH=7.2). FACS was used to determine the quality of the nuclei.

### Mitochondrial studies

Tissue was harvested from adult *Mus* and *Acomys* as approved by the University of Michigan animal protocol PRO00010623. Mice were injected intraperitonially 7 minutes prior to tissue harvesting with ketamine (100 mg/kg) and dexmedetomidine (0.5 mg/kg) as anesthesia followed by heparin (Hospira; 1000 U/kg) at 5 minutes prior to tissue harvesting. The aorta was cannulated while in the thoracic cavity and perfused with an ice cold cardioplegia solution (25 mM KCl [P4504, Sigma], 100 mM NaCl [S9888, Sigma], 10 mM Dextrose [D9559, Sigma], 25 mM MOPS [M1254, Sigma] 1 mM EGTA [E4378, Sigma]), pH 7.2) for 5 minutes on iceThe apex of the heart was used for the citrate synthase assay and the remainder of the heart was used in the purification of mitochondria.

#### Mitochondrial purification

Mitochondria were purified as previously described [21]. A portion of the LV and septum was minced in a beaker on ice for 3 minutes and subsequently was homogenized using a glass dounce homogenizer at 4°C for 3 minutes in 10 mL of isolation buffer (200 mM Mannitol [M9647, Sigma], 60 mM sucrose [S7903, Sigma], 5 mM KH2PO4 [P5379, Sigma], 5 mM MOPS [M1254, Sigma], 1 mM EGTA [E4378, Sigma], pH 7.2) and 3 mg of proteinase (P8038, Sigma). After 3 minutes, 25 mL of isolation buffer with 0.1% w/w BSA (A6003, Sigma) and protease inhibitor (539134, Calbiochem, Darmstadt, Germany) was added and the homogenate was centrifuged at 8,000 x g for 10 min at 4°C. The pellet was resuspended in 25 mL of isolation buffer with BSA and centrifuged at 8,000 x g for 10 min at 4°C. The resulting pellet was resuspended in isolation buffer with BSA and centrifuged at 700 x g for 10 min at 4°C. The supernatant was collected and centrifuged at 8,000 x g for 10 min. The mitochondrial pellet was resuspended in a minimal amount of isolation buffer with BSA.

#### Citrate synthase assay

Mitochondrial protein concentration was measured using the Quick Start Bradford assay (#5000205, BioRad Laboratories, Hercules, CA) and the Quick Start Bovine Gamma Globulin Standard Set (#5000209, BioRad). The citrate synthase assay was previously published by Oroboros. Briefly, 2 µg of purified mitochondria were placed in a solution containing 100 mM Tris pH 8.1 (T1503, Sigma), 0.25% triton X-100 (T8532, Sigma), 0.31 mM acetyl CoA (A2181, Sigma; and 10101907001, Roche, Basel Switzerland), 0.1 mM DTNB (D218200, Sigma), and 0.5 mM oxaloacetate (04126, Sigma) at 30°C. Immediately, absorbance was measured at a wavelength of 412 nm, and the rate was determined and used to calculate citrate synthase activity.

To measure cardiac tissue citrate synthase activity, 30 mg of fresh heart tissue is minced in 100 mM Tris pH 7. Once minced, the tissue is homogenized in a glass dounce at 4°C 3 times for 30 seconds every 2 minutes. The homogenate is diluted 1:15 in 1% triton X-100 and 100 mM Tris pH 7.0. The diluted homogenate is incubated on ice while vortexing every 3 minutes for 30 minutes total. The homogenate is further diluted 1:40 in the 1% triton X-100 and 100 mM Tris pH 7.0. The citrate synthase activity assay is performed as stated above, where instead of 2 µg of protein, 20 µL is used.

#### Mitochondrial respiration

Mitochondrial respiration was measured from purified mitochondria using a high-resolution Oxygraph O2k (Oroboros Instruments GmbH; Innsbruck, Austria). Following the protocol previously published [21], each chamber contains 2 mL of respiration buffer (90 mM KCl [P4504, Sigma], 50 mM MOPS [M1254, Sigma], 1 mM EGTA [E4378, Sigma], and 0.1% w/w BSA [A6003, Sigma], pH 7.2) with 5 mM KH2PO4 [P5379, Sigma] and is set to 37°C while stirring at 750 rpm. Mitochondria was added so that each chamber had 0.54 U of citrate synthase activity to establish the baseline respiration in state 1. For state 2, the substrate of choice was added to the chambers at the following concentrations: 1) 5 mM sodium pyruvate (P8574, Sigma), 2 mM malate (M1000, Sigma), and 5 mM NaCl (S9888, Sigma), or 2) 0.020 mM palmitoyl-L-carnitine (P1645, Sigma), 2 mM malate, and 10 mM NaCl. After mitochondria and substrate were added, 1 mM K-ADP (A5285, Sigma) was added to each chamber for state 3.

#### NAD(P)H Autofluorescence

Mitochondria were suspended in 1mL respiration buffer in 24-well black assay plates (4titude 4ti-0262) at the same concentration as in the respirometry experiments determined by the CS assay. NAD(P)H autofluorescence was measured from the top of each well at 37°C in a modular multi-mode microplate reader with an automated injection system (BioTek Synergy H1, Agilent). The excitation and emission wavelengths were set to 340nm and 460nm, respectively. The gain and read height were automatically adjusted by the Gen5 3.11 software to optimize fluorescence measurements of mitochondria suspended in respiration buffer in the presence of 2.5mM KH2PO4, 5mM NaCl, 0.5mM sodium pyruvate, 0.25mM potassium malate pH 7.0, and 10*μ*M rotenone (Sigma R-8875). The gain was set to 134 and the read height 6.75mm. Only 4 wells were used at once to increase the number of data point collected during each run. For all runs, fluorescence in the 4 wells was measured every 6 seconds with double orbital mixing in between measurements. Experiments were run in duplicate and the raw fluorescence data were averaged before analysis. For each experimental run, depending on the substrates provided to the mitochondria, NaCl was added to achieve a final sodium concentration of 10 mM. For example, 9.5mM NaCl was added to runs with 0.5mM sodium pyruvate, while no NaCl was added when 5mM disodium succinate was present. Mitochondria were added to 1mL of RB containing NaCl, if needed, and KH2PO4, mixed manually, bubbles were removed, and fluorescence was measured over 2:10 minutes. Substrate(s) were automatically injected, and the plate was mixed for 5 seconds before measuring state 2 for 2:30 minutes. After the automatic addition of potassium ADP and 5 seconds of mixing, state 3 was measured for 5 minutes. To measure the fully reduced state, 10*μ*M of rotenone was added manually, the plate was mixed for 5 seconds, and fluorescence was measured for 3 minutes. In runs where pyruvate and malate were not already present, 0.5mM sodium pyruvate and 0.25 mM potassium malate pH 7.0 were added with rotenone to ensure a fully reduced state.

### Single-nucleus RNA sequencing data analysis

Libraries were sequenced using the Novaseq 6000. Resulting fastqs were processed through CellRanger v2.2.0 (https://support.10xgenomics.com/single-cell-geneexpression/software/pipelines/latest/what-is-cell-ranger) using either the mm10 genome (for *Mus*) or an annotated *Acomys* genome to obtain a gene expression matrix. To help with downstream analyses, *Acomys* genes were annotated with their mouse ortholog name if there was a one-to-one match, i.e., one gene from *Mus* matched one gene in *Acomys*.

Analyses were performed using Seurat v3 [22, 23] and custom R scripts generally following the PBMC tutorial (https://satijalab.org/seurat/articles/pbmc3k_tutorial.html) and the Integration tutorial (https://satijalab.org/seurat/articles/integration_introduction.html). Cells were filtered for cells containing over 500 genes, mitochondrial content below 20%, and hemoglobin below 2.5%. The mitochondrial and hemoglobin filtering was only performed in *Mus* samples because of missing annotation in *Acomys*. Cells were included in the analysis if they had at least 750 features. Only one read per cell was needed for a gene to be counted as expressed. The expression depth for each cell was normalized to 10000 genes per cell and log transformed [24]. The data was scaled to reduce the effect of sequencing depth of each sample by regressing out this factor. When merging all data sets together, dimension reduction batch correction was performed by merging the data sets using 2000 anchor genes common to all data sets found within the FindIntegrationAnchors function and a final merging using the IntegrateData function. Dimension reduction was performed using UMAP (Uniform Manifold Approximation and Projection) using genes common to both species. Cell clusters were determined by the Louvain algorithm at a clustering resolution of 0.5. Twenty PCA components were used for clustering and UMAP. Tentative cluster annotation was done with scCATCh [25] with the tissue set to “Blood”, “Heart” or “Heart muscle” and the p-value threshold set to 0.05. CM clusters were positively identified based on the expression of Myh6, and Tnnt2. Any non-CM clusters were removed from further analyses.

Cluster markers were identified using the ‘FindConservedMarkers’ function in Seurat with the MAST statistical test [26]. Differential expression was performed using the ‘FindMarkers’ function also with the MAST statistical test. Genes were submitted for statistical analysis if they had at least 0.1 of cells expressed that gene. Differentially expressed genes (DEGs) were identified based on a pval <0.05, and a log2 fold change (log2FC) cutoff of 1.5.

**Monocle3:** To identify which CM population gives rise to mCM2 in *Mus* and aCM3 in *Acomys* after injury, we performed pseudotime trajectory analysis using the Monocle3 v1.3.1 package, as described in the documentation here (https://cole-trapnell-lab.github.io/monocle3/docs/introduction/) [27, 28]. Cells from mCM2 and aCM3 cluster were used as the input expression matrix for trajectory analysis at Day1, 3, or 7 as compared to their respective sham controls. Cells were then ordered along pseudotime with CM1 population as the “root cells”, and colored by their original identities. Differential gene expression analysis was performed using the graph-autocorrelation analysis to find genes that vary over a trajectory.

**GATOM:** The network analysis to elucidate significant interactions was performed using GATOM [29]. Overall, it consists of two major steps: 1) Creating a metabolic network using either genes and/or metabolites, 2) Identifying a subnetwork that encompasses the most significant changes.

To run the gene-based analysis, we converted the snRNAseq data into a “pseudo-bulk” dataset using the ‘aggregateExpression’ function available in the Seurat v5.1.0 package. Next, DESeq2 was run to obtain a list of genes, and their baseMean values. A pval <0.05 and log2FC cutoff of 1.5 was used to identify the DEGs. The list of DEGs were uploaded to GATOM to construct network using a network topology of atoms in the KEGG database.

To run the metabolite-based analysis, differentially expressed metabolites were used as inputs for network construction in GATOM using a network topology of metabolites in the KEGG database. Student’s T-test was used to identify differentially expressed metabolites between the comparison groups.

**Data and code availability:** The cell Ranger Single-Cell Software Suit is found here https://support.10xgenomics.com/single-cell-geneexpression/software/pipelines/latest/what-is-cell-ranger

**Seurat guided clustering can be found here:** https://satijalab.org/seurat/articles/pbmc3k_tutorial.html

**Monocle 3 package was used as described in the tutorials here:** https://cole-trapnell-lab.github.io/monocle3/

**The shiny GATOM app is available at** https://artyomovlab.wustl.edu/shiny/gatom/

### Statistics

Values are expressed as mean ± standard error of mean (s.e.m). We used student T-test, one-way-ANOVA, repeated measures ANOVA and mixed-effects ANOVA with Tukey’s corrections to compare data across species as appropriate. The mixed-effects model integrates analysis that uses the available data to estimate model parameters. Animals with partial data contribute to the estimation of some model parameters but not others. It is important to note that relying solely on repeated-measures ANOVA will limit the sample size to only animals that survived for the entire duration of the study and could lead to selection bias as we are only selecting a subgroup of animals. Throughout the manuscript, analyses that compared species and time but did not include paired samples such as data in Figure 4G-H were analyzed using 2-way independent ANOVA. Sample sizes, statistical tests and P values are indicated in the figures or figure legends. Animal numbers are presented as a range based on the number of animals included in each analysis. These numbers are included in the figure legends. Throughout the analyses, a P value < 0.05 was considered statistically significant.

Metabolomics raw data was processed using the Agilent software (MassHunter Qual and ProFinder). Data analysis was performed with Agilent MassProfiler Pro package using recursive analysis workflow.

## Results

### Single-cell RNA sequencing reveals distinct transcriptional responses to myocardial infarction in regenerative versus non-regenerative cardiomyocytes

Enhanced cardiac repair in *Acomys* [7] suggests a novel cardiomyocyte (CM) phenotype compared to non-regenerative mammals. Single nucleus RNA sequencing (snRNA-seq) enabled high-resolution dissection of CM heterogeneity and dynamic injury responses in the regenerative *Acomys* heart compared to the non-regenerative *Mus* heart. Unbiased clustering of >50,000 CM nuclei per species uncovered 4 distinct populations in *Acomys* (aCM1-4) and 5 in *Mus* (mCM1-5) under homeostatic conditions (**Figure 1A-B**).

**Figure 1.**
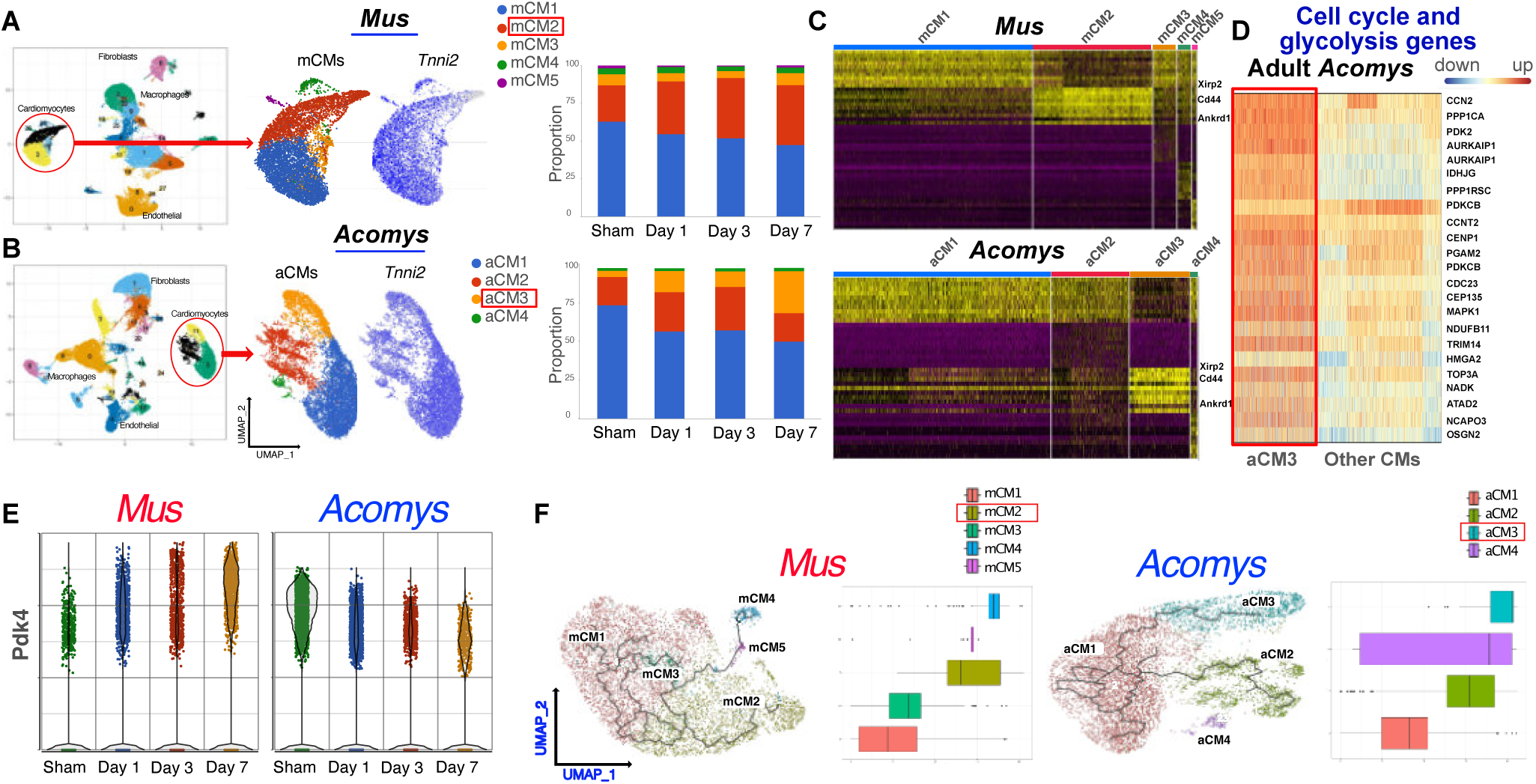
Adult *Acomys* CMs exhibit unique features and divergent MI responses compared to *Mus*. Single-nucleus RNA sequencing with unbiased clustering of all time points reveals distinct cardiomyocyte clusters in *Mus* (mCMs) **(A)** and *Acomys* (aCMs) **(B)** hearts after myocardial infarction. UMAP visualization and proportions of snRNA-seq defined CM cluster populations and days 1-7 post-MI in *Mus* and *Acomys* show cardiomyocyte (CM) clusters in *Acomys* (aCM) and *Mus* (mCM) at homeostasis and D1, D3, and D7 post-MI. Feature plots showing cluster-specific expression of CM structural protein Tnnt2 in CM clusters. The dynamics of cardiomyocyte clusters after MI demonstrate a highly dynamic cluster in *Mus* (mCM2) and *Acomys* (aCM3). (**C**) Heatmaps display the top marker genes for the *Mus* and *Acomys* CMs with mCM2 cluster in *Mus* highly expressing Xirp2, Ankrd1, while aCM3 cluster in *Acomys* highly expressing *Xirp2*, *Cd44*. (**D**) Heat maps of selected cell cycle and glycolysis genes demonstrate that aCM3 has significantly higher proliferation and cell cycle gene expression than other CM clusters. These data indicate sustained reliance on fatty acid oxidation regardless of MI duration or snRNA-seq cluster in *Mus* but shift towards glycolysis in *Acomys*. **(E)** Violin plot of *Pdk4* gene expression in *Acomys* and *Mus* at baseline and after MI in *Acomys* and *Mus* demonstrates the higher expression of *Pdk4* in *Acomys*. **(F)** Pseudotime analysis reconstructing activation of gene expression programs along inferred developmental trajectories and branchpoint decisions. This analysis suggests that mCMs transition to become predominantly mCM2 while *Acomys* predominantly evolve into aCM3. Nuclei were pooled from 3-5 animals/group/time point.

Following myocardial infarction, major modulation of CM cluster proportions occurred over 7 days after MI (**Figure 1**). While mCM1 nuclei decreased only slightly in abundance, the mCM2 population dramatically expanded from 24% in sham-operated animals to 40% by day 7 post-MI. Concomitantly, top mCM2 markers *Ankrd1*, *Xirp2*, and *Nppb* transcript levels were increasingly upregulated over this timeframe. In contrast, the analogous aCM3 cluster increased from 4.2% to 10.4% of total CM nuclei by day 7 post-MI in *Acomys*, albeit with less pronounced induction of markers like *Xirp2* and *Cd44*.

Importantly, we observed higher expression of cell cycle and glycolysis genes in aCM3 compared to other CM clusters in *Acomys* (**Figure 1D and Suppl. Figures 1 and 2**). In support of these observations, the expression level of *Pdk4*, a major switch between fatty acid oxidation and glycolysis in cardiomyocytes, decreased post-MI in *Acomys* while was upregulated post-MI in *Mus* (**Figure 1E**).

Pseudotime trajectory analysis positions single CM nuclei along a continuum of inferred developmental states, allowing the reconstruction of lineage relationships. Applied to the *Mus* MI time course data, this approach placed mCM2 downstream of a branchpoint decision, supporting origination from pre-existing CM populations rather than the emergence of a distinct progenitor (**Figure 1F**). The mCM2-traversing trajectory showed marked upregulation of gene programs linked to pathological hypertrophy, apoptosis, mitochondrial dysfunction, arrhythmogenesis, and suppressed angiogenesis over 7 days post-MI. Targets induced include fetal genes *Myh7*, *Acta1*, and *Nppb*, apoptotic mediator *Casp3*, fibrotic driver *Tgfb1*, ion channels *Scn5a* and *Cacna1c*, and fatty acid translocase *Cd36.* Concurrently, pro-angiogenic factors like *Vegfa* and *Pecam1* were increasingly downregulated along the pseudotime path. In contrast, mCM1 maintained a consistent transcriptional signature post-MI with minimal deviation. Our pseudotime trajectory analysis indicated a rapid emergence of the mCM2 and aCM3 populations from pre-existing CM clusters in *Mus* and *Acomys*, respectively. This emergence is likely due to transcriptional remodeling in response to injury, supporting the snRNA-seq findings and underscoring the metabolic shifts occurring in these CM populations.

In summary, snRNA-seq analysis identified important inherent differences between cardiomyocytes in *Acomys* and *Mus*. Post-MI, our data suggests different trajectories and responses to ischemic injury.

### Cardiomyocyte metabolic shifts are linked to survival Post-MI

Metabolic adaptability is particularly important during periods of stress or injury, such as MI, when the energy demands of the heart are significantly increased [30]. In our snRNAseq studies of CM clusters in both species, we observed a higher utilization of different metabolic substrates, including glycolysis in *Acomys*, while *Mus* relied heavily on fatty acid oxidation (**Suppl. Figure 1 and 2**). This finding is consistent with previous research, which reported that metabolic changes associated with cardiomyocyte dedifferentiation enable adult mammalian cardiac regeneration [15, 31]. When we focused our analysis on mCM2 and aCM3 clusters, we observed a similar pattern regardless of MI status (**Figure 2A**). No activation of glycolytic metabolism was detected over 7 days post-MI in *Mus*, even in the expanding mCM2 population. This persistent catabolic inflexibility may indicate that *Mus* CMs have limited capability to adapt after MI, which may contribute to impaired ischemic recovery capacity.

**Figure 2.**
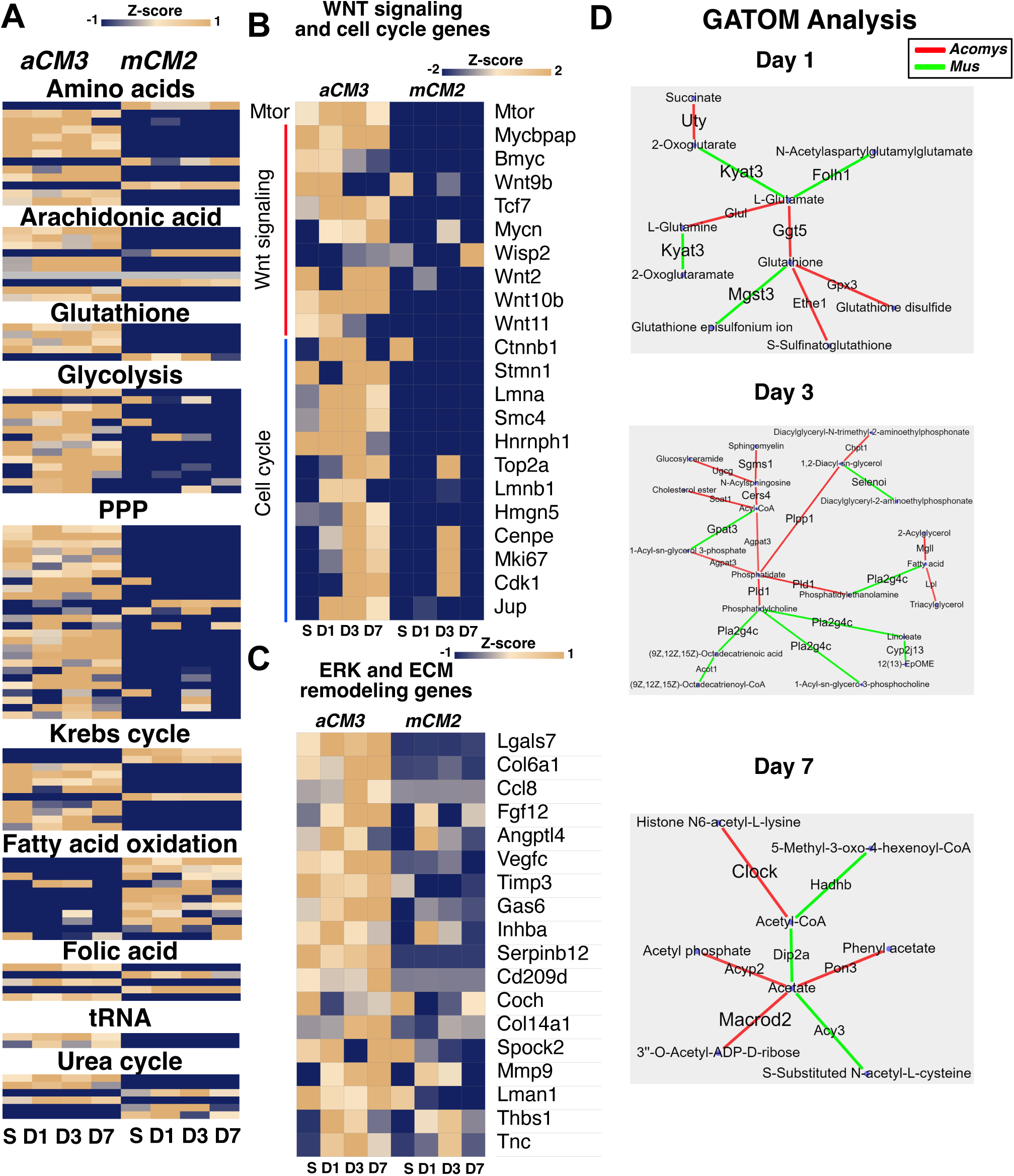
Characterization of *Acomys* and *Mus* metabolic response to MI reveals divergent phenotypes. Transcriptomic data using snRNAseq in sham (S), and days 1,3, and 7 (D1, D3, and D7) post-MI, show higher expression of glycolysis, PPP, amino acid catabolism, and glutathione metabolism genes in aCM3 compared to mCM2. Conversely, genes involved in fatty acid oxidation are higher in mCM2 (nuclei were collected from 3-5 animals/group/time point). ***Acomys* aCM3 exhibits a higher expression of *Wnt* signaling and proliferation genes compared to *Mus* mCM2.** snRNAseq studies in CM clusters, in sham (S), and days 1,3, and 7 (D1, D3, and D7) post-MI, reveal higher expression of mTOR, Myc] and Wnt signaling genes, and cell cycle/proliferation genes after MI in *Acomys*, particularly at day 3 post-MI (nuclei were pooled from 3-5 animals/group/time point). C. **Network analysis by GATOM of metabolic gene expression in aCM3 vs. mCM2 on days 1, 3, and 7 post-MI.** aCM3 demonstrated higher activity of antioxidant and detoxification genes compared to *Mus* (green lines). On the other hand, mCM2 demonstrated higher levels of FA oxidation and TCA cycle activity (red lines).

In addition to the unique glycolytic characteristics, *Acomys* CMs demonstrate higher levels of activity in multiple biosynthetic pathways, such as the pentose phosphate pathway (PPP) and glutathione metabolism (**Suppl. Figure 1**). Biosynthetic pathways, such as the PPP, could influence CM proliferation by several means. PPP plays an important role in the de novo synthesis of nucleotides, which are required for RNA and DNA synthesis and repair [32]. Additionally, pentose phosphate and serine biosynthesis pathways are important for NADPH regeneration and NAD+ synthesis [32], which are important for sustaining adequate levels of NAD(H) for dehydrogenase reactions [15, 33]. Indeed, our snRNAseq data indicate that *Acomys* aCM3 demonstrates a high activity of the PPP, which has been shown to activate the mTOR pathway [34], with subsequent activation of Wnt signaling and CM proliferation genes, particularly at day 3 post-MI (**Figure 2B**). A combined analysis of all CM clusters showed similar patterns (**Suppl. Figure 1**). These findings offer insights into the unique metabolic characteristics of CMs in adult *Acomys*. These observations were evident both in sham-operated animals and after MI with adaptive increase in some of the glycolysis and PPP genes. Furthermore, the expression of Redox genes was higher in *Acomys* CMs, which supports cell proliferation and produces ROS scavengers (**Figure 3**).

**Figure 3.**
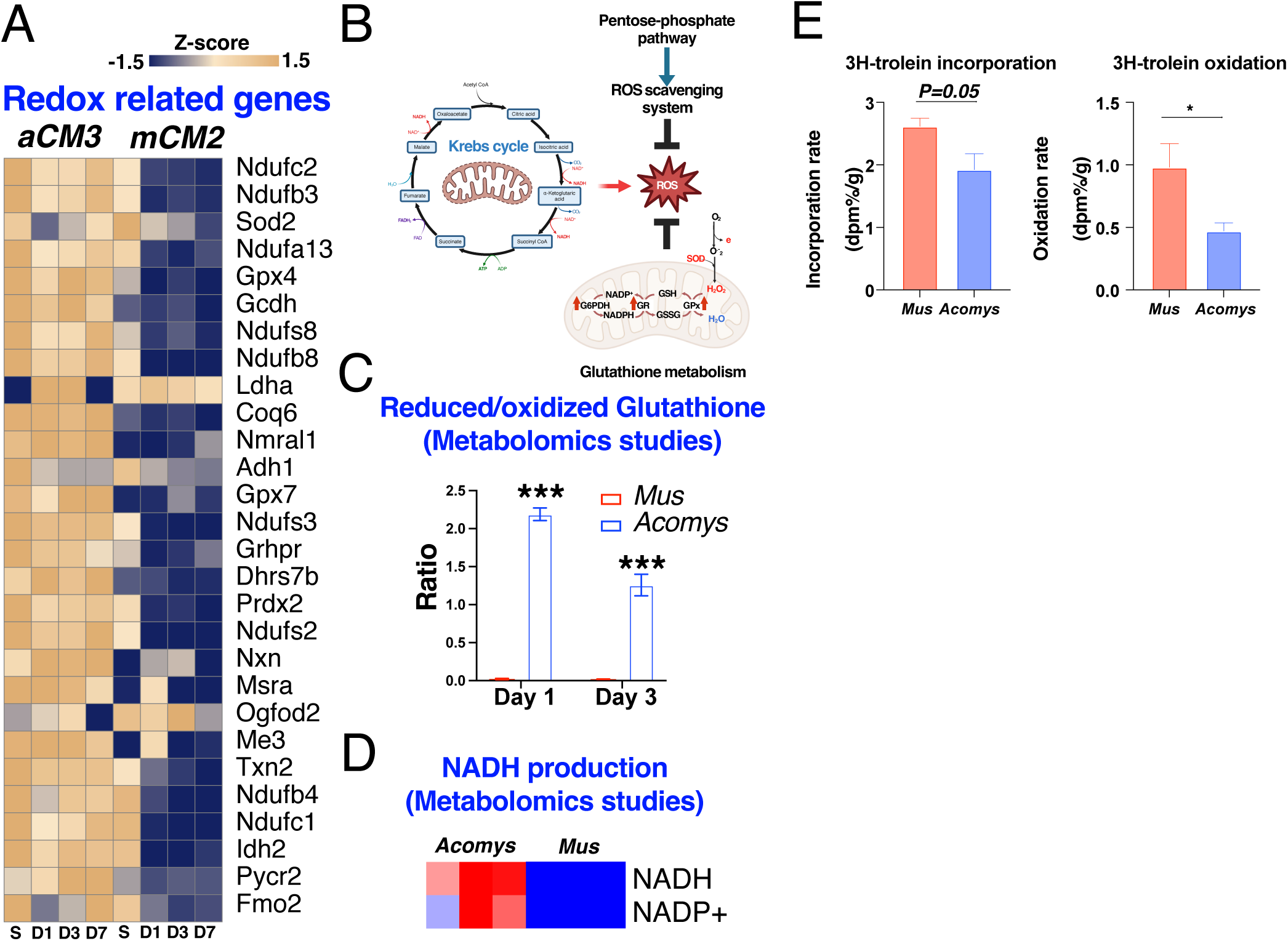
*Acomys* adopt unique metabolic pathways to maintain Redox balance after MI. *A*. snRNA-Seq data demonstrates increased expression of enzymes involved in redox balance in aCM3 compared to mCM2, sham (S), and days 1,3, and 7 (D1, D3, and D7) post-MI. **B.** schematic of ROS production and clearance in Acomys. **Panels C & D** depict targeted metabolomics analysis of heart tissue at 1- and 3-days post-MI, showing the higher ratio of reduced/oxidized glutathione and NADH/NADP+ in Acomys, reflecting the more efficient neutralization of ROS (3-5 animals/group/timepoint for snRNAseq and 3 animals/group/timepoint for metabolomics. **E. Total 3H radioactivity in cardiac tissue at the end of the 3H-triolein injection experiment.** Radioactivity is calculated as a percentage of input and normalized against tissue weight. The data suggests lower free fatty acid incorporation and oxidation in *Acomys* compared to *Mus*. (3-5 animals/group, \**P*<0.05 compared to *Mus*).

Further analysis of the metabolic state of aCM3 vs. mCM2 was performed using GATOM [29], a method of unbiasedly analyzing and integrating ‘omics data for understanding metabolic regulation in cellular processes. This analysis revealed distinct differences in the TCA cycle, amino acid metabolism, and antioxidant defense systems. *Mus* hearts exhibited higher flux through the TCA cycle, as evidenced by increased conversion of 2-oxoglutarate to L-glutamate by Oxoglutarate Dehydrogenase L (*Ogdhl*), and potentially a more oxidized state (red nodes, **Figure 2C**). In contrast, *Acomys* hearts upregulated antioxidant defenses as indicated by higher activity of the glutathione synthesis pathway (green nodes, **Figure 2C**). Furthermore, *Acomys* hearts maintains an elevated NADP(H)/NADP ratio at baseline and has an increased capacity to utilize carbohydrates, even after MI (**Figure 3**).

These findings suggest that enhanced carbohydrate metabolism and generation of PPP precursors are critical inherent characteristics, allowing *Acomys* CMs to maintain redox balance by upregulating the machinery needed to neutralize ROS. These unique changes allow *Acomys* CMs to proliferate and withstand ischemic injury, as we have previously shown [7].

### Metabolic adaptations in adult *Acomys* cardiomyocytes promote ROS neutralization

The cardiac *Mus*cle relies on oxidative phosphorylation for energy production, which generates ROS. Under normal conditions, endogenous antioxidants regulate ROS levels, which are crucial for cardiomyocyte proliferation, function, and survival. However, during ischemic injury, excessive ROS production overwhelms the antioxidant capacity, leading to oxidative stress [9], damaging cellular components, impairing contractile function, and causing cell death. Relying on glycolysis helps reduce ROS production, which can cause DNA damage and accelerate cell cycle arrest in neonatal CMs [10].

To understand cardiac metabolic responses to myocardial infarction, we conducted targeted metabolomics using LC/MS mass spectrometry of the entire heart in sham-operated and injured animals. our metabolomic data corroborates the transcriptomic data and shows higher glutathione synthase activity in *Acomys* hearts favors increased capacity for ROS scavenging as shown in the ratio of reduced to oxidized glutathione levels. Baseline metabolic profiling of *Acomys* and *Mus* hearts revealed significant differences in their metabolic signatures. *Acomys* exhibit distinct metabolic profiles with higher impacts in pathways such as riboflavin metabolism, amino sugar and nucleotide sugar metabolism, and the pentose phosphate pathway compared to *Mus*. This suggests that *Acomys* have a unique baseline metabolic state that might contribute to their cardiomyocyte survival capabilities.

Notably, pathways such as alanine, aspartate, and glutamate metabolism, as well as arginine biosynthesis, were upregulated in *Acomys* (**Figure 4A-C**).

**Figure 4.**
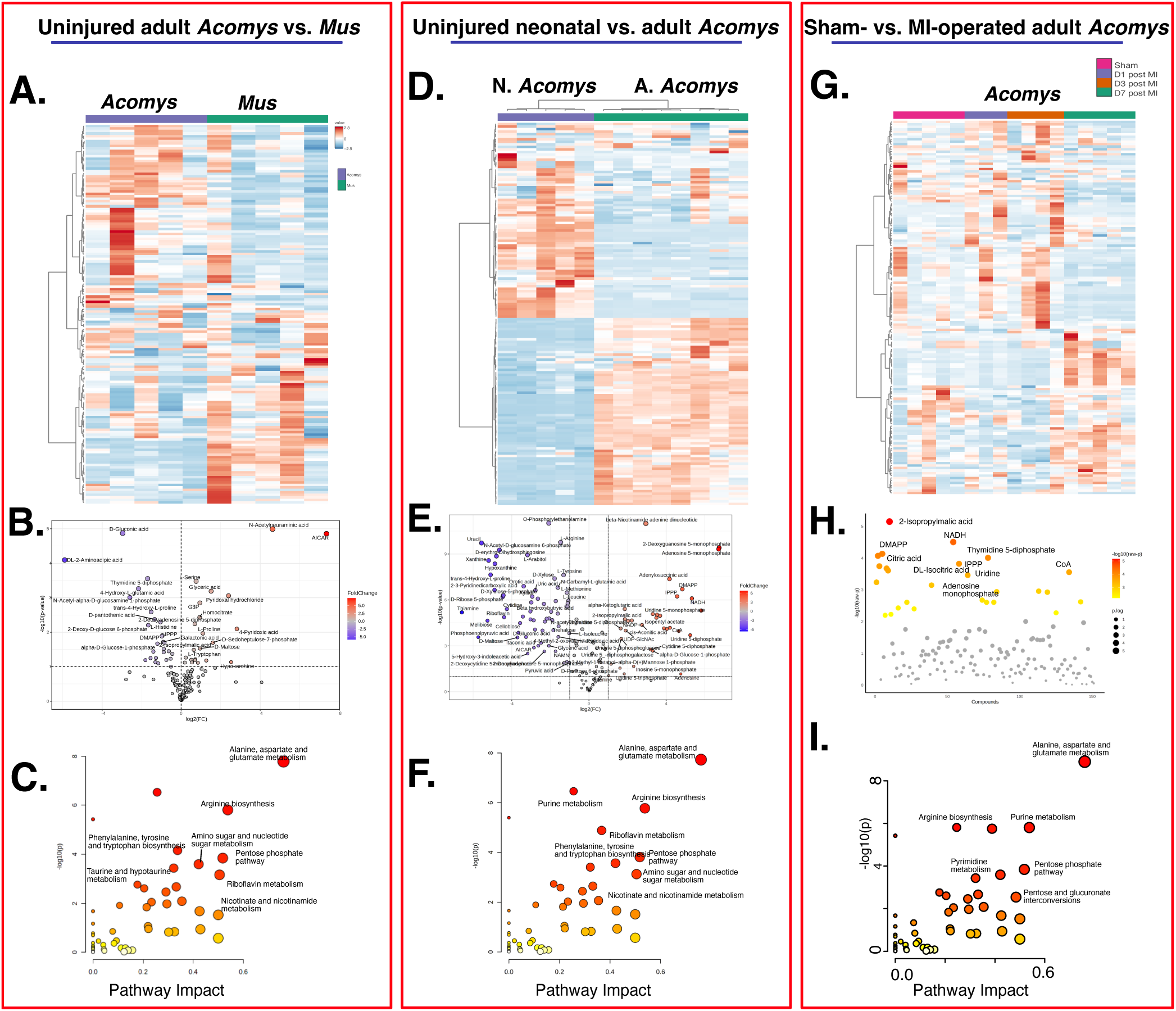
Metabolomic Profiling of *Acomys* and *Mus* Rodents in Baseline and Post-Myocardial Infarction Conditions. This figure presents a comprehensive metabolomic analysis comparing *Acomys* and *Mus* across various physiological states, including neonatal, adult, sham-operated, and post-myocardial infarction conditions. Panels **A, D, and G** display heat maps of metabolite levels for baseline comparisons between adult and neonatal *Acomys* (A. *Acomys* and N. *Acomys*) and *Mus* (A), neonatal vs. adult *Acomys*, and adult *Acomys* at days 1, 3, and 7 after MI, sham-operated animals. Color intensity indicates relative normalized metabolite abundance (red for higher levels, blue for lower levels). The data highlights the dynamic metabolic differences between *Acomys* and *Mus* at baseline and the upregulation of these metabolic changes in response to myocardial infarction. Panels **B, E, and H** illustrate Volcano plots of the most abundant metabolites in neonatal and adult *Acomys* at baseline and after myocardial infarction. Panels **C, F, and I** further explore pathway analysis of metabolic pathways, showing differences between *Acomys* and *Mus* and between neonatal and adult *Acomys*. Each point represents a metabolic pathway, with the x-axis showing pathway impact and the y-axis showing -log(p-value) of the enrichment analysis. Color coding from yellow to red indicates increasing pathway impact. This data suggests upregulation of amino acid metabolism, arginine biosynthesis, purine metabolism, and pentose phosphate pathway activity in *Acomys* compared to *Mus*. These metabolic pathways are further upregulated in *Acomys* after myocardial infarction in response to tissue injury, offering insights into the metabolic adaptations and responses to MI in both species. The data suggest significant species-specific and condition-specific metabolic alterations, which may affect understanding the resilience and recovery mechanisms in myocardial infarction (N = 3-6 animals/group/time point).

When compared to neonatal *Acomys*, adult *Acomys* hearts demonstrate a pronounced shift in metabolic activity related to purine metabolism, riboflavin metabolism, and nicotinate and nicotinamide metabolism (**Figure 4D-F**). These changes suggest an adaptive metabolic response aimed at enhancing energy production and reducing oxidative stress.

After myocardial infarction, *Acomys* further adapt to ischemic stress with significant shifts in metabolic profiles over the first week after the ischemic insult. Our data suggest increased abundance of 2-isopropylmalic acid, NADH, and thymidine 5’-diphosphate, which are implicated in energy production and cellular repair. Pathway impact analysis indicates that pathways like alanine, aspartate, and glutamate metabolism, as well as purine metabolism, are significantly impacted post-injury, suggesting their crucial role in the metabolic adaptation and survival of cardiomyocytes in *Acomys* (**Figure 4G-I**).

Taken together, the results indicate that *Acomys* mice exhibit a distinct metabolic profile that supports enhanced cardiomyocyte survival. The upregulation of pathways involved in energy production, antioxidant defense, and cellular repair processes at baseline, coupled with adaptive metabolic changes post-injury, underscores the metabolic resilience of *Acomys* cardiomyocytes. These findings provide valuable insights into the metabolic mechanisms underlying cardiomyocyte survival and highlight the potential of *Acomys* as a model for studying cardiac metabolism and ischemic resilience.

### *Acomys* demonstrate higher cardiomyocyte proliferation after MI compared to *Mus*

We observed higher levels of cell cycle genes in *Acomys* CMs. To further explore the level of CM proliferation in *Acomys* CMs, we conducted immunohistochemistry (IHC) studies using antibodies against EdU in animals treated with EdU at 10 mg/kg for 14 days starting the day of MI. Our IHC analyses demonstrated a significantly higher number of EdU+ CMs in *Acomys* compared to *Mus* (**Figure 5 A&B**). We then conducted studies to quantify cytokinesis events using a recently developed cell fate mapping and highly specific cytokinesis/M-phase reporter [17]. Aurora kinase B (AurKb) is one of the major protein kinases that ensures the proper execution and fidelity of mitosis and is only expressed for a short time during the M-phase/cytokinesis process, localizing to the central spindle during anaphase and in the midbody during cytokinesis [18]. To quantify cytokinesis events, we recently developed a cytokinesis/M-phase reporter using an *AurKb* promoter-based GFP reporter and cloned it into a non-integrating lentivirus to drive the permanent expression of GFP protein specifically in CMs using the TNNT promotor [17]. We have validated that this reporter can track *Acomys* neonatal CM division *in vitro* and labels proliferating CMs *in vivo* after MI in *Mus* and *Acomys* **(Figure 5C-F).** Animals received intramyocardial Injection of the AurKb-GFP-IL lentiviral reporter system injected in the peri-infarct region at 4.5×10^7^ transducing units (TU) immediately following LAD ligation, and we assessed the rate of GFP+ cells at 14 dpi to provide definitive evidence regarding CM proliferation across species. We observed significantly higher numbers of GFP+ CMs in the peri-infarct region in *Acomys* compared to *Mus* (**Figure 5G&H**). This was associated with a smaller size of CMs in *Acomys* compared to *Mus* (**Figure 5I&J**). Overall, these findings corroborate our snRNA-seq studies and suggest that the unique metabolic phenotype in *Acomys* CMs could be associated with their enhanced ischemic resilience.

**Figure 5.**
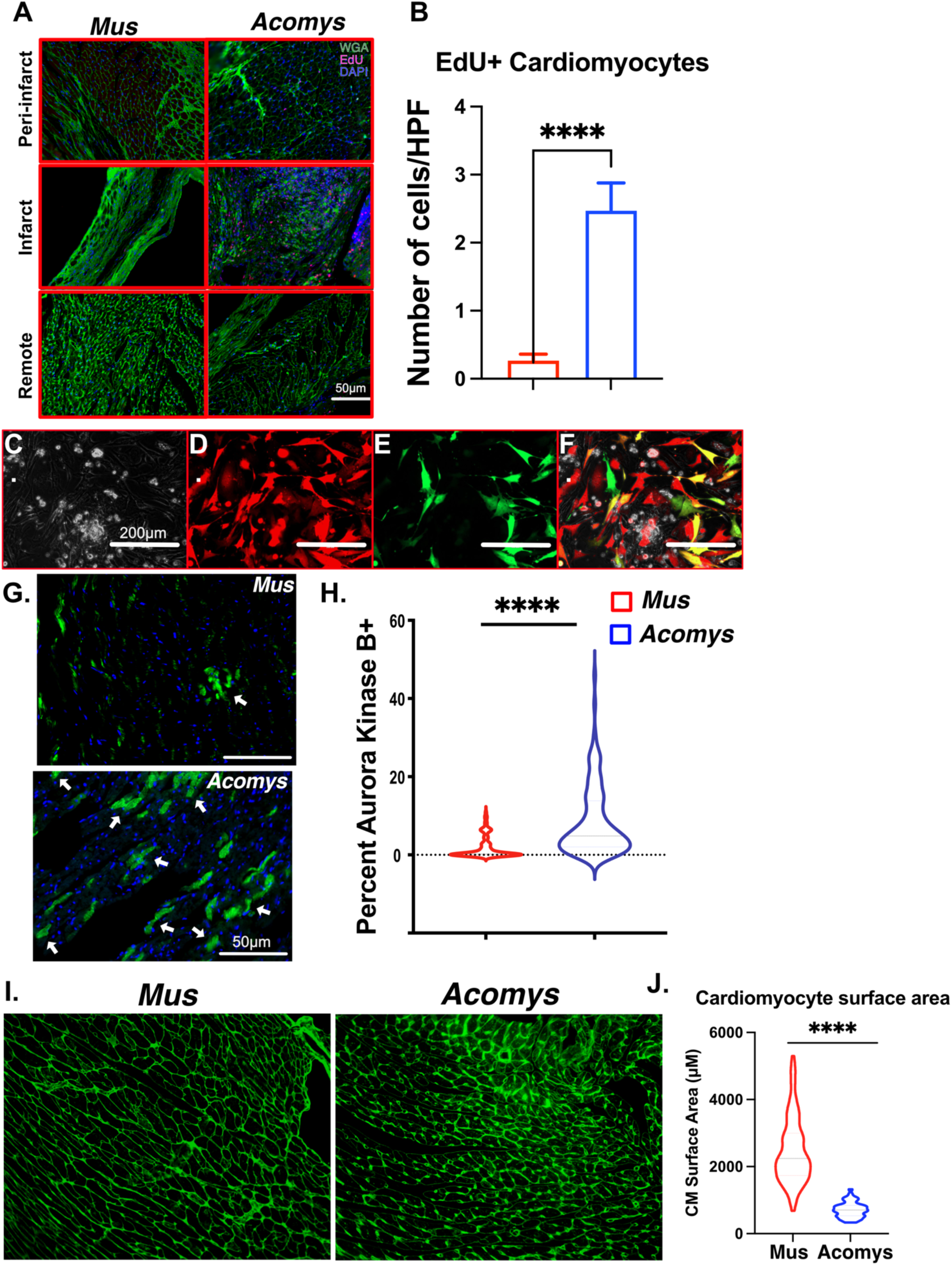
*Acomys* exhibit cardiomyocyte proliferation after ischemic injury. *Mus* and *Acomys* were injected with EdU at 80m mg/kg for 14 days, starting the day of MI. 14 days after MI, animals were sacrificed, and heart sections were stained against EdU and the cardiac-specific protein TnnT. **A & B** show a significantly higher number of EdU+ CMs in the peri-infarct region in *Acomys* (3-4 animal/group, ****P<0.0001, scale bars = 50 μm). **C-F** *Acomys* neonatal CMs were isolated as previously described [19] and cultured on laminin (Gibco) coated plates. Cells were then transfected with IL DR and IL Aurora cre constructs (2×10^5^ TU/μl). Panels C-F represent Brightfield, DsRed, GFP, and merged images showing successful transfection and labeling of cardiomyocytes 3 days after transfection (4 replicates scale bars = 200 μm). **G & H** show successfully labeled proliferating cells in the peri-infract zone 3 days after MI and transfection with a significantly higher number of GFP+ cells in *Acomys*. Panel **I** shows Wheat Germ Agglutinin staining of cardiac sections to visualize the CM cell border. Panel **J** demonstrates the quantification of CM size in *Acomys* and *Mus* under physiological conditions.

### *Acomys* hearts exhibit metabolic phenotypes similar to neonatal mice

Unlike other mammals, *Acomys* demonstrate superior ischemic tolerance and cytoprotection following MI, which is associated with smaller scar size. Our preliminary studies show that *Acomys* CMs, similar to proliferating cells, utilize a diverse range of fuel substrates, including glucose, purines, amino acids, and glutathione. This metabolic adaptation maintains the balance between energy production and ROS generation, which is critical for CM survival and cell-cycle progression. Mitochondrial respiratory studies using the Oxygraph O2k (Oroboros Instruments) revealed that *Acomys* mitochondria have an enhanced capacity to oxidize carbohydrates relative to fatty acids and generate a greater NADP(H)/NADP ratio compared to *Mus* (**Fig. 6**). *Acomys* maintain a higher pyruvate to palmitoyl-L-carnitine (PLC) ratio even in adulthood compared to *Mus* in sham and MI-operated animals (**Fig. 6A-C**), explaining their higher cardiac reduced/oxidized glutathione cardiac tissue (**Figure 3**). Mitochondrial content in *Acomys* hearts was significantly lower than in *Mus*, regardless of age and injury status, reflecting their reduced reliance on oxidative phosphorylation (**Fig. 6D**). This data suggests that *Acomys* deploy additional antioxidant mechanisms that reduce CM apoptosis, synergistically improving cardiac repair beyond cell proliferation. We then conducted radiolabeled in vivo lipid chase studies at the Michigan Mouse Metabolic Phenotyping Center (Mi-MMPC). Uninjured semi-fasted *Acomys and Mus* underwent a 3 H-triolein chase study to assess FA oxidation, as previously published [35]. *Mus* appears to have a higher rate of fatty acid oxidation, suggesting a mitochondrial substrate preference for fatty acids over carbohydrates for energy production. The difference in triolein uptake was primarily the result of a reduced level of incorporation (**Figure 4**). These differences in fatty acid metabolism could have implications on the energy balance and ROS production between *Mus* and *Acomys*, as observed in our mass spectrometry studies (**Figure 3**).

**Figure. 6.**
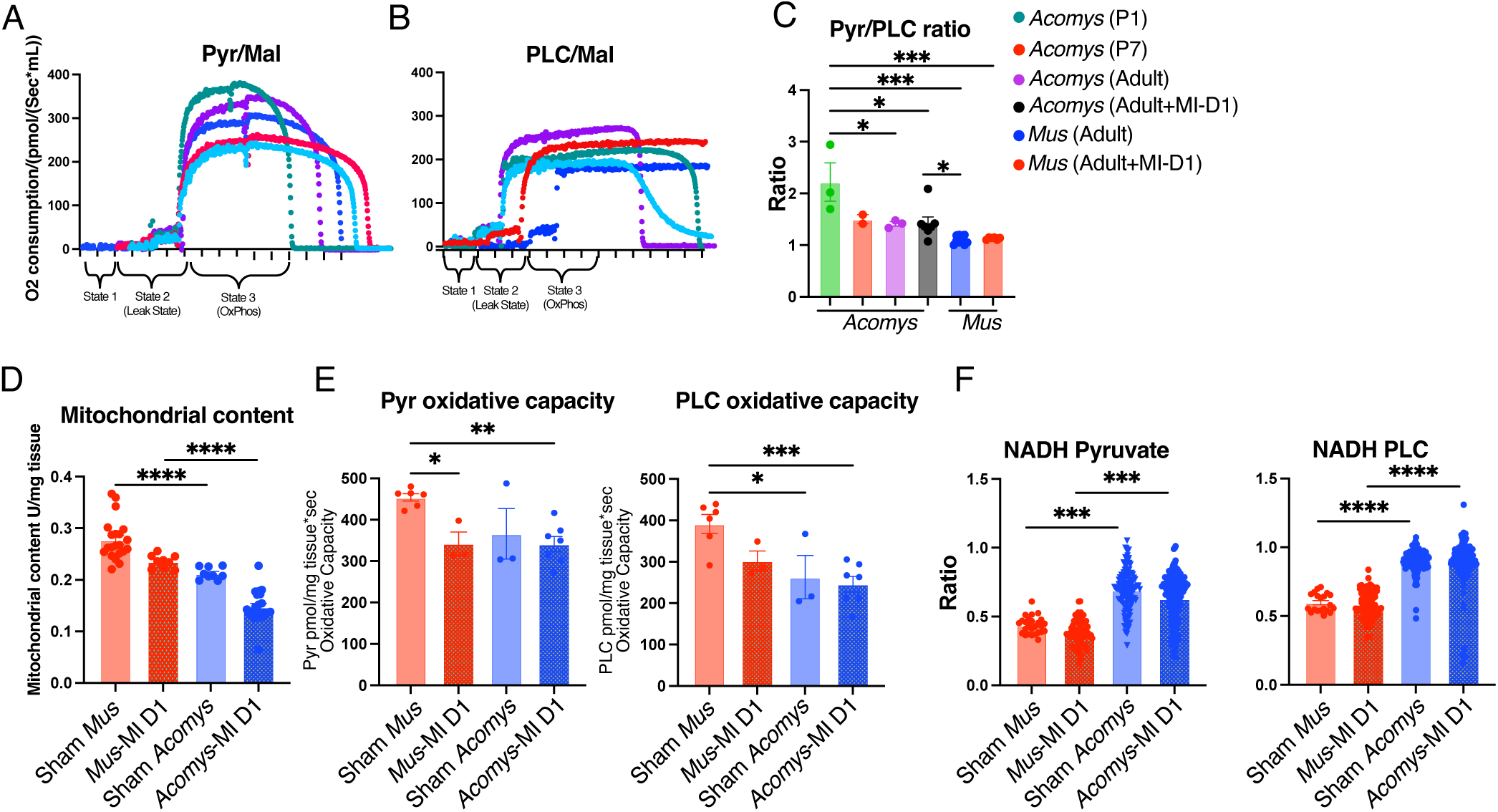
Mitochondrial studies corroborate the transcriptomic and metabolomic data and demonstrate higher glycolysis and NADH production in *Acomys*. Mitochondrial respiration rates in isolated mitochondria from *Acomys* and *Mus* hearts in sham-operated mice showed higher pyruvate utilization (glycolysis) in *Acomys* **(A)** and comparable palmitoyl-L-carnitine (PLC) utilization **(B)**; which resulted in higher pyruvate/PLC ratio in *Acomys* compared to *Mus* **(C)**. The pyruvate/PLC ratio was highest in neonatal *Acomys* and decreased in adult *Acomys*. However, adult *Acomys* had a significantly higher pyruvate/PLC ratio than adult *Mus*. Additionally, *Acomys* had lower mitochondrial content per mg of heart tissue than *Mus* **(D)**. **(E)** Mitochondrial studies show reduced oxidative capacity with both pyruvate and PLC in *Acomys* compared to *Mus*. Post myocardial infarction (MI); there is a reduction in oxidative capacity in *Mus* but not in *Acomys*. **(F)** At baseline and post-MI, *Acomys*’ mitochondria produced higher rates of NADH with both pyruvate and PLC utilization. Values in means ± S.E.M. (n=3-6 animals/group, \**P*<0.05, \*\*\**P*<0.001, \*\*\*\**P*<0.0001 compared to *Mus*).

## Discussion

Cardiomyocyte metabolism plays a crucial role in their proliferation, survival, and response to ischemic injury. Although existing literature has established the role of metabolism in the regenerative capacity of neonatal cardiomyocytes, there is a paucity of data on these pathways in adult mammals [2, 8, 10]. The present study addresses several limitations of previous research on cardiac regeneration. By comparing CM responses to myocardial infarction in both *Mus* (common mice) and *Acomys* (spiny mice), this study provides a comprehensive understanding of the potential variability in these responses across non-regenerative and regenerative species. Furthermore, it offers a detailed analysis of the metabolic shifts in cardiomyocytes following MI, revealing a shift towards glycolysis in *Acomys*, in contrast to the reliance on fatty acid oxidation in *Mus*. Lastly, the study identifies specific pathways associated with reparative and maladaptive responses to MI, with *Acomys* showing upregulation of pathways related to cytoskeleton organization, adrenergic signaling, oxidative stress resistance, angiogenesis, and metabolic pathways, while *Mus* exhibited upregulation of pathways associated with cellular senescence, hypertrophy, and arrhythmogenic right ventricular cardiomyopathy. Our findings provide novel insights into the synergistic metabolic adaptations of ischemic resilience and recovery in adult CMs and provide important therapeutic targets for ischemic heart disease..

Our approach of examining transcriptomic and functional inherent and adaptive metabolic response to ischemia provides a powerful approach to elucidate mechanisms of ischemic resilience in *Acomys*. Our finding that *Acomys* CMs possess unique inherent characteristics that increase glycolytic flux while avoiding maladaptive hypertrophy gives insights into how ischemic recovery is regulated. Identifying strategies to unlock similar plasticity in adult human CM represents a promising therapeutic direction. Akin to proliferating CMs, our transcriptomic and functional studies revealed that *Acomys* hearts downregulate fatty acid oxidation (oxidative machinery) and augment anabolic pathways such as glucose-derived carbon allocation into the NAD+, pentose phosphate, glutathione synthesis, and other ancillary biosynthetic pathways. These pathways have been shown to promote CM proliferation, reduce ROS production, and enhance their detoxification with increased synthesis and salvage of adenine nucleotides [15]. This unique set of synergistic metabolic adaptations in *Acomys* presents a unique and perhaps superior model for studying the protection of the adult heart during ischemia and could inform the design of future successful therapies.

Our previously published baseline comparisons of CM characteristics between *Acomys* and *Mus* identified several unique features in *Acomys* typically associated with immature or neonatal *Mus* CMs, such as smaller size, higher percentage of mononucleated CMs, presence of T-type calcium channels, and reduced t-tubule density and organization [7, 16]. This immature phenotype has been linked to higher regenerative potential in other models [36–42]. In fact, *Acomys* CMs exhibit a metabolic profile that favors glycolysis over fatty acid oxidation, similar to neonatal cardiomyocytes [10, 11], a metabolic phenotype that is associated with reduced reactive oxygen species (ROS) production and survival during hypoxia. When subjected to myocardial infarction, *Acomys* demonstrated remarkable ischemic tolerance and myocardial preservation, resulting in reduced adverse remodeling, smaller scar size, and improved survival compared to *Mus* strains.

ROS homeostasis is crucial in cardiomyocyte proliferation and survival following ischemic injury. Excessive ROS production during ischemia-reperfusion can lead to oxidative stress, DNA damage, and cell death [9]. In contrast, maintaining a balanced ROS level is essential for cardiomyocyte proliferation and heart regeneration [8, 10]. Our transcriptomic in CMs and functional data in the entire heart suggest that *Acomys* deploy coordinated and synergistic metabolic pathways that reduce ROS production and enhance their detoxification. These pathways are inherent to *Acomys* cardiomyocytes with additional adaptive response to ischemia. The higher glycolytic activity and reduced reliance on fatty acid oxidation in *Acomys* cardiomyocytes may contribute to lower ROS generation compared to adult *Mus* cardiomyocytes [14,15]. Furthermore, the upregulation of antioxidant genes such as Prdx1, Sod1, Sod2, and G6pd in *Acomys* CMs indicates an enhanced capacity to neutralize ROS and maintain redox balance. The higher activation state of the pentose phosphate pathway and glutathione metabolism in *Acomys* cardiomyocytes may also protect against oxidative stress by generating NADPH and glutathione, essential cofactors for ROS scavenging enzymes [15]. The unique ROS homeostasis in *Acomys* cardiomyocytes may confer a significant advantage in their survival following ischemic injury. By maintaining a balanced ROS level, *Acomys* cardiomyocytes can reduce the detrimental effects of oxidative stress on cell cycle progression and viability. This is consistent with studies showing that reducing ROS production or enhancing antioxidant defense can promote cardiomyocyte proliferation and improve cardiac recovery post-MI [6,8,12].

In conclusion, our study establishes the unique metabolic phenotype underlying the ischemic resilience and post-MI recovery in *Acomys* cardiomyocytes, such as enhanced ROS homeostasis and a shift toward glycolysis and ancillary biosynthetic pathways. These pathways favor cell survival, proliferation, and tissue regeneration. These findings provide valuable insights into the metabolic mechanisms underlying cardiac repair and highlight potential therapeutic targets for improving outcomes in ischemic heart disease. Future studies should explore whether pharmacological or genetic modulation of these pathways can enhance repair following ischemic injury.

## Acknowledgments

We acknowledge support from the Bioinformatics Core of the University of Michigan Medical School’s Biomedical Research Core Facilities (RRID:SCR_019168).

