## Supplemental Figures for "Inherent Metabolic Adaptations in Adult Spiny Mouse (*Acomys*) Cardiomyocytes Facilitate Enhanced Cardiac Recovery Following Myocardial Infarction"

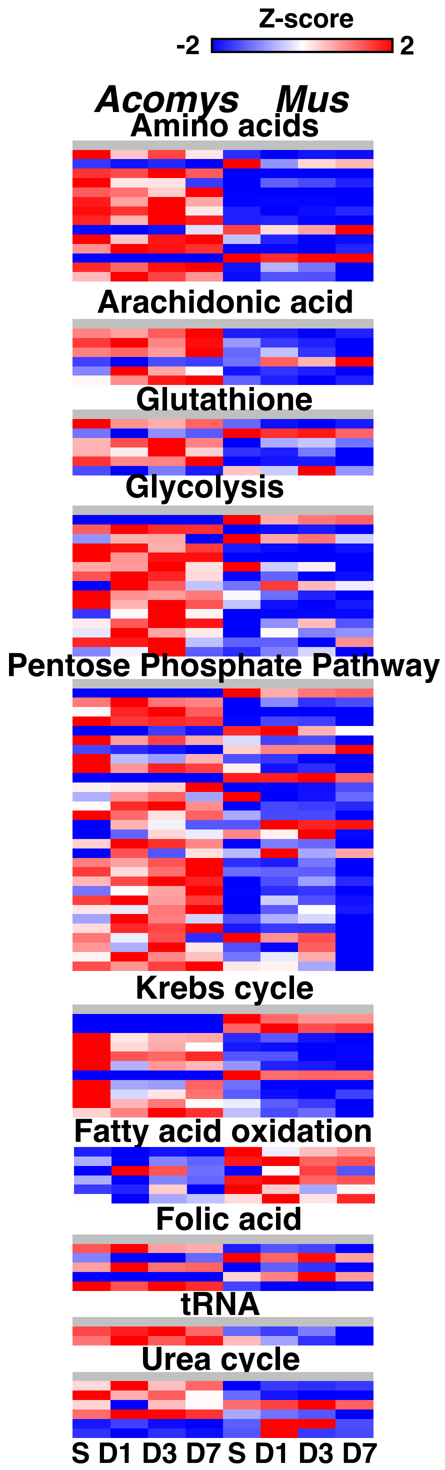


**Suppl. Figure 1. Characterization of *Acomys* and *Mus* metabolic response to MI reveals divergent phenotypes**. Transcriptomic data using snRNAseq in sham (S), and days 1,3, and 7 (D1, D3, and D7) post-MI, show higher expression of glycolysis, PPP, amino acid catabolism, and glutathione metabolism genes in *Acomys* vs. *Mus* CMs. Conversely, genes involved in fatty acid oxidation are higher in <us compared to *Acomys* (nuclei were collected from 3-5 animals/group/time point).

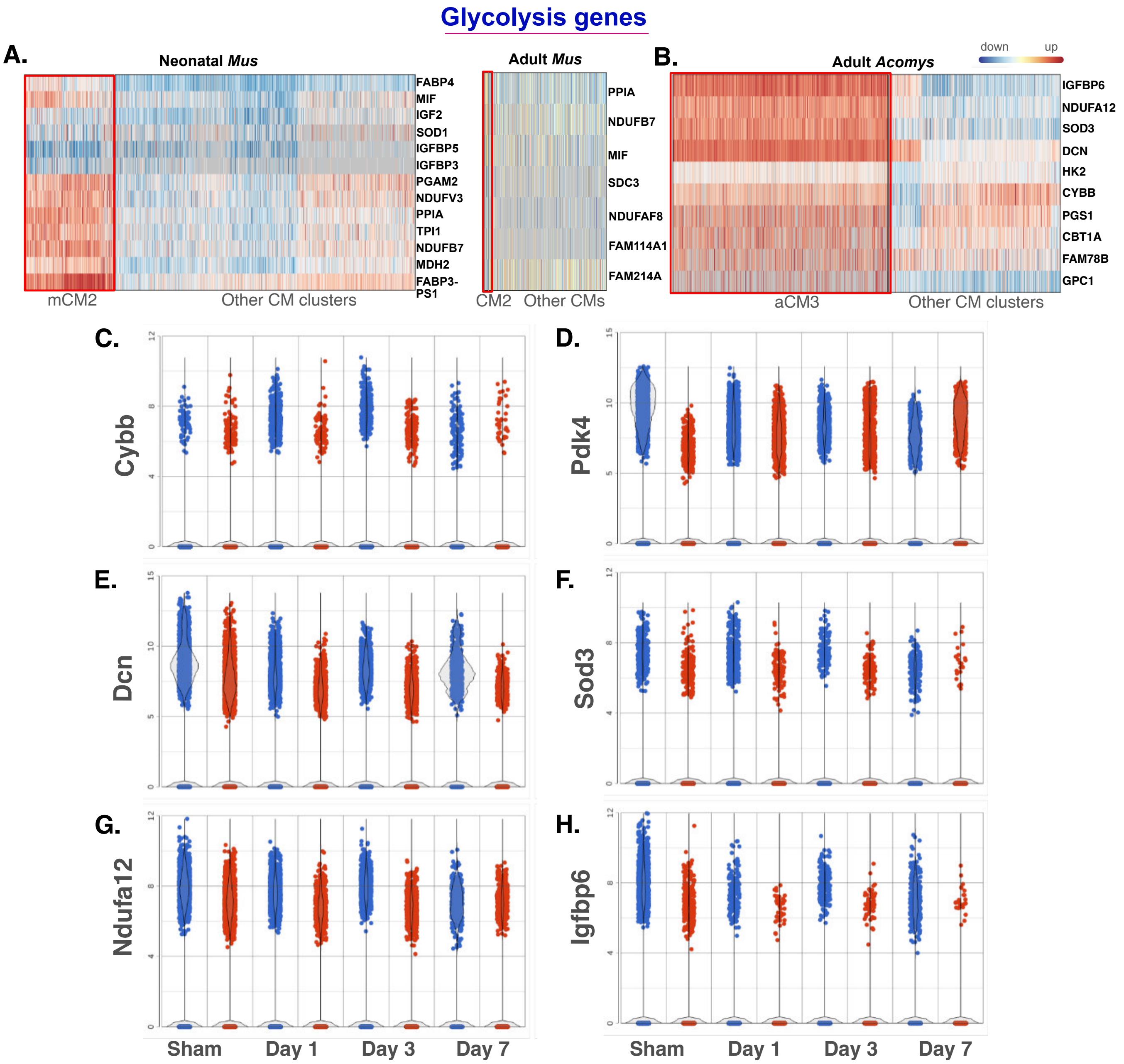


**Supplemental Figure 2. *Acomys* cardiomyocyte cluster 3 resembles neonatal proliferating cardiomyocytes.** Gene expression changes in neonatal mouse and adult *Mus* and *Acomys* cardiomyocytes. The heatmap at the top shows expression levels of selected glycolysis genes at various time points in proliferative neonatal mouse cardiomyocytes (mCM2, **A**) compared to other neonatal cardiomyocyte clusters. **B.** gene expression of glycolysis-related genes in aCM3 compared to other *Acomys* CM clusters. **C-H.** Volcano plots display expression changes over time (Sham, Day 1, Day 3, Day 7) for genes involved in glycolysis (*Cybb, Pdk4, Dcn, Sod3, Ndufa12,* and *Igfbp6*) in *Acomys* CMs compared to *Mus* (n=3-5 animals/group/time point).
